# Reevaluating the Association Between Epstein-Barr Virus (EBV) and Breast Cancer in the United States

**DOI:** 10.1101/2024.11.28.625954

**Authors:** Clarence C. Hu, Devanish N. Kamtam, Juan J. Cardona

## Abstract

The World Health Organization estimates 9.9% of cancers are attributable to viruses. Notably, human papillomavirus causes roughly 90% of cervical cancers, while Epstein-Barr virus (EBV) is linked to nearly 10% of gastric carcinomas. Regarding breast cancer, the association with EBV is inconclusive. While studies in some nations report an association, those in the United States largely do not. We reviewed studies from 2003 to 2023 and identified seven that analyzed EBV association with breast cancer in American patients. We observed a potential risk of not investigating novel EBV variants. Detection protocols utilized only lymphoma-derived strains, despite the current knowledge suggesting that genotype variation can influence pathogenic potential and cell tropism. Certain EBV strains, for instance, may preferentially infect epithelial cells and increase the risk of nasopharyngeal carcinoma (NPC) by up to 11 times. Stated simply, the optimal EBV detection protocol for breast cancer cells may differ from lymphoma cells. Reliance on lymphoma-derived strains assumes a level of sequence conservation among EBV genomes. Mounting evidence demonstrates greater variation than previously believed, especially in key coding and non-coding regions. Our analysis reveals that 5/7 (71%) studies used at least one assay sequence that did not exactly match more than 50% of EBV genomes in NCBI GenBank. Moreover, 98% of these GenBank entries became available after assay sequences were selected. Overall, it is possible the current understanding may be incomplete. Should breast cancer mirror gastric carcinoma and exhibit EBV influence in certain subtypes, these insights could enable targeted therapies and screening programs.

**Objectives:** This study examines potential limitations of prior investigations into the association between Epstein-Barr Virus (EBV) and breast cancer in the United States. Specifically, our aims are to:

1. **Assess the cellular origin and pathogenicity of EBV strains employed in detection protocols.** This objective stems from the background section’s discussion on EBV genotype variations and their potential influence on tissue tropism and pathogenic mechanisms.
2. **Evaluate the sequence similarity between assay sequences and available EBV genomic data.** This objective addresses the concern raised in the background section regarding the potential for newfound sequence variation among EBV strains and the implications for accurate detection.
3. **Determine the extent to which detection protocols incorporate the latest EBV genomic data.**

## Background

### Viral Association With Cancer

The World Health Organization (WHO) estimates that 15.4% and 9.9% of all cancers are attributable to infectious organisms and viruses, respectively(1). An extensive study investigating the association between cancers and viruses using whole genome sequencing led by the Pan-Cancer Analysis of Whole Genomes (PCAWG) consortium identified 16% of the cases to be associated with viruses(2).

Cancers with established viral etiology or strong association with viruses include:

- Cervical cancer(3,4)
- Burkitt lymphoma (BL)(5)
- Hodgkin lymphoma(6)
- Gastric carcinoma(7)
- Kaposi’s sarcoma(8)
- Nasopharyngeal carcinoma (NPC)(9–11)
- NK/T-cell lymphomas(6)
- Head and neck squamous cell carcinoma (HNSCC)(12)
- Hepatocellular carcinoma (HCC)(13)

### Viral Mechanisms of Action in Cancer

Viruses may promote multiple stages of carcinogenesis, including initiation, progression, and therapeutic resistance.

Viruses are known to influence key proteins, pathways, and chromosomal sites implicated in tumorigenesis, including:

- MYC(14–17)
- TP53(15,18,19)
- PD-L1(20–25)
- BRCA1(26,27)
- EGFR(28,29)
- CDK6(8,30)
- STAT3(31)
- MTHFD2(17)
- MLL(32)
- LARG(32)
- PI3K-Akt(33)
- JAK-STAT(34)

In addition, viruses may also induce MDR1 overexpression and reduce treatment efficacy with only a few infected cells.(35)

### EBV Overview

EBV is a double-stranded DNA virus from the Herpesviridae family that is formally classified as human gammaherpesvirus 4. It can be transmitted asymptomatically for weeks via common body fluids like breast milk, saliva, and blood. EBV infects over 90% of individuals worldwide, usually asymptomatically. EBV infection typically occurs early in life as approximately 50% of children carry EBV by age 10 and 80% by age 18.(36,37) While EBV infection classically presents as infectious mononucleosis (mono), it is also known to be causally associated with multiple sclerosis.(38) EBV establishes lifelong persistence by tethering to host chromosomes and downregulating immune activity to escape immune surveillance.(39) While EBV predominantly colonizes B lymphocytes, it has also been detected in epithelial cells and T lymphocytes.

### EBV Association With Malignancies

EBV is classified as a class 1 carcinogen and is strongly associated with several cancers, including Burkitt lymphoma, Hodgkin lymphoma, gastric carcinoma, NK/T-cell lymphomas, and NPC.(40)

### EBV Association With Breast Cancer

The association between EBV and breast cancer is inconclusive.

Studies from China, India, southern Europe, and a few African nations have demonstrated a higher prevalence of EBV in breast cancer samples than in normal breast tissue.(41–43) Notably, some studies report an association between EBV and triple-negative breast cancer (TNBC), an aggressive phenotype disproportionately affecting younger patients, with one study identifying 36% prevalence of EBV in TNBC samples.(44,45) In the USA, however, studies have largely demonstrated no association between EBV association and breast cancer.

Moreover, mice models have also demonstrated that EBV infection of mammary epithelial cells promotes malignant transformation and initiates uncontrolled growth.(46)

### EBV Life Cycle and Genome

The EBV genome consists of 170k-180k DNA base pairs, encoding over 80 proteins and 40 non-coding RNAs.(47)

Similar to other herpesviruses, EBV is characterized by a latent-lytic life cycle. During the lytic stage, the virus is extremely immunogenic, producing the broad array of gene products required for viral replication and infection. Conversely, the latent stage expresses a sparse set of gene products and is typically undetectable to the immune system.

EBV is believed to exist primarily in the latent stage, which comprises four sub-stages, or types, marked by disparate protein and RNA expression: 0, I, II, and III.

Only three gene products, all of which are non-coding RNAs, are expressed in every latency sub-stage: EBV-encoded RNA 1 (EBER1), EBV-encoded RNA 2 (EBER2), and BamHI-A rightward transcript (BART). EBER1 and EBER2 are abundantly expressed during latency. Among their many functions is the ability to suppress or confer resistance to the host immune response mediated by interferon (IFN)-α and T helper 1 (Th1) cells. EBERs bind to protein kinase R (PKR) and inhibit PKR phosphorylation in order to evade IFN-α-induced apoptosis.(48,49)

The protein expressed in the most stages, Epstein-Barr virus nuclear antigen 1 (EBNA1), is silent during type 0 latency.

EBV genotypes are classified as type 1 or type 2 (type A or type B, respectively). Type 1 EBV strains are prevalent worldwide, whereas type 2 strains are more common in tropical regions like Papua New Guinea and sub-Saharan Africa.(50)

### Incomplete EBV Genome Understanding

Although the oncogenic potential of EBV was discovered in 1964, the impact of its genomic diversity on oncogenic profiling and clinical outcomes remains incompletely understood, particularly when compared to the advanced typing systems used for human papillomavirus (HPV).(51–56)

These knowledge gaps stem from technical challenges and a historical shortage of data. A large genome and many repetitive sequences make complete-genome sequencing for EBV costly and time-consuming. For comparison, the EBV genome is 170k-180k base pairs while HPV is about 8k base pairs.(57)

Prior to 2006, only four complete EBV genomes were available and only one of type 2 before 2015.(58–61) Recent advances in technology, however, have augmented these numbers. As of May 2024, the National Center for Biotechnology Information (NCBI) GenBank lists 512 complete EBV genomes, and 1,269 partial or nearly complete genomes.

### Emerging Non-coding and Coding Genomic Variability

While variations in a few genes and non-coding regions have been documented, many regions of the EBV genome remain understudied. Several studies have attempted to correlate genetic variations with disease prevalence, but encountered challenges due to the lack of sequence-specific clinicopathological data. Furthermore, recombination events among distinct EBV strains may introduce confounding factors that are difficult to disentangle.(59,60)

The recent increase in genomic data has revealed more pronounced diversity in coding and non-coding regions than previously believed. Notably, commonly used detection targets belong to regions with emerging variability, including EBER1, EBER2, EBNA1, BART, BZLF1, and LMP1.(52,56,62–64,64–69) Palser et al. (2015) have also reported that the single nucleotide polymorphism (SNP) density varies substantially across all known open reading frames and notably, is highest in latency-associated genes.(59)

### Genomic Variability Impacts Pathogenic Potential and Cell Tropism

DNA viruses such as HPV and EBV, although more stable than RNA viruses, nonetheless undergo mutations that may alter pathogenic potential and cell tropism.

### Pathogenic Potential

The vast majority of HPV genotypes are non-tumorigenic. Among over 150 HPV genotypes, only seven drive approximately 90% of cervical cancer cases while two alone, HPV 16 and HPV 18, account for 70%.(70)

EBV subtypes may also exhibit similar differences in oncogenic potential between different subtypes. Type 1 and type 2 EBV strains initiate cellular transformation and proliferation with varying degrees of efficiency and consequently also report different malignancy rates.(65,71–73) Xu et al. (2019) observed an 11-fold increase in the risk of NPC progression due to EBV isolates with a specific variant in the BALF2 gene, namely BALF2_CCT.(74)

Other studies also demonstrate the impact of small genomic variations on lytic replication, progression, and etiopathogenesis of NPC and other cancers.(53,75–79) Even SNPs may enhance oncogenic potential, illustrated by a single nucleotide substitution amplifying lytic reactivation.(80) Besides pathogenic potential, SNPs may also correlate with geographical differences. Patients with NPC in Japan demonstrated a unique EBV subtype with single nucleotide variations that varied from the ones prevalent in NPC-endemic regions like southern China.(66) Importantly, the pathogenicity of a strain may correlate with cell infectivity and specificity. The M81 strain, isolated from a patient with NPC, preferentially infects epithelial cells and demonstrates a higher incidence of NPC.(78)

EBER polymorphisms strongly associate with high-risk variants of NPC and appear more often in type 1 strains.(62,63) Different EBER subtypes also correspond to different rates of leukemia and myelodysplastic syndrome (MDS).(52) Different EBV strains may rely on different genes for pathogenesis. For instance, the EBER2 mechanism, which accelerates cell division by upregulating UCH-L1 deubiquitinase and indirectly overexpressing cyclin B1 and Aurora kinase B, is more crucial for cellular transformation in the M81 strain than in others. Different cell types may also exhibit different levels of EBER2 dependent proliferation as Li et al. (2021) demonstrated with B cells and epithelial cells. Significantly, EBER2 may be indispensable for Burkitt lymphoma pathogenesis since every oncogene except EBNA1 typically remains silent.(80) All told, these polymorphisms are noteworthy because detection protocols may rely on dated EBER gene sequences, despite indications of greater heterogeneity than previously understood.

### Cell Tropism

Cell/tissue tropism reflects the ability of a pathogen to selectively infect specific organs or a group of organs.(81)

HPV genotypes show distinct cell tropism, preferentially infecting squamous and glandular epithelium.

EBV primarily infects B cells and epithelial cells, and less frequently NK cells and T cells.(82)

Different EBV strains may exhibit enhanced tropism for epithelial cells over B cells or vice versa. Epithelial-tropism of EBV, and glandular tropism in particular, remains under-explored and incompletely understood.(51,78)

### EBV Reference Genomes and Cell Lines

The NCBI lists two official reference genomes for EBV: B95-8 for type I and AG876 for type II.(83–85) The Raji strain, another widely used reference genome, was isolated from the Raji cell line.(59,86,87) Two common EBV cell lines are Raji and Namalwa.

While the B95-8 strain was isolated from monkey lymphocytes infected with EBV from a patient with infectious mononucleosis, the AG876 strain was isolated from patients in Africa with Burkitt lymphoma.(88–92) The Raji strain originated from an 11-year-old male with Burkitt lymphoma.(86)

The Raji and Namalwa cell lines were also isolated from patients in Africa with Burkitt lymphoma.(86,89–92)

The first complete EBV genome derived from a patient with carcinoma, GD1 (GenBank accession AY961628), was published in 2006 but derived from saliva donated by a male Cantonese patient in China with NPC and not derived from carcinoma cells.

### Potential EBV Detection Challenges in Adenocarcinomas

HPV and EBV oncogenic models suggest that viral detection in adenocarcinomas may require protocols accounting for low copy numbers, single nucleotide mismatches, and strain multiplicity. Moreover, Arbach et al. (2006) concluded that EBV genomes may distribute unevenly in breast tumors, with one area containing high copy numbers while another yields low copy numbers.(35) While the data does not claim these factors are unique to adenocarcinomas, it does suggest the need for higher specificity and sensitivity. Studies found that viral DNA in glandular cells may present in low copy numbers and trigger false negatives even with single nucleotide mismatches.(78,93,94) Furthermore, individuals may carry multiple EBV strains, which may suggest that isolates derived from saliva and non-tumor cells may not represent isolates in tumor cells.(58,95) This underscores the need to restrict reference strains to those collected from cancer cells, avoiding strains such as GD1 that are isolated from saliva and non-tumor cells.

### RNA Integrity Number (RIN)

RNA Integrity Number (RIN) measures RNA integrity and ranges from 1 to 10. Scores of 1 indicate completely degraded RNA while 10 indicates intact RNA. Although a RIN of 5 is generally acceptable for routine use, scores of 8 or higher are recommended for the most sensitive applications.(96,97) Low RIN values may compromise accuracy and reflect poor tissue quality or management.

Given the importance of accurate detection protocols, sequence specificity, and biomolecule integrity in determining the association between EBV and breast cancer, our study aimed to reevaluate the methodologies and findings of prior research from American studies conducted in the past 20 years. We sought to identify potential biases and gaps in the current understanding of this association and their implications on the reported association between EBV and breast cancer.

## Materials and Methods

### Literature Search Methodology

We searched PubMed for studies spanning the last 20 years due to significant advancements in Epstein-Barr virus (EBV) research and availability of novel EBV genomic data over this time period.

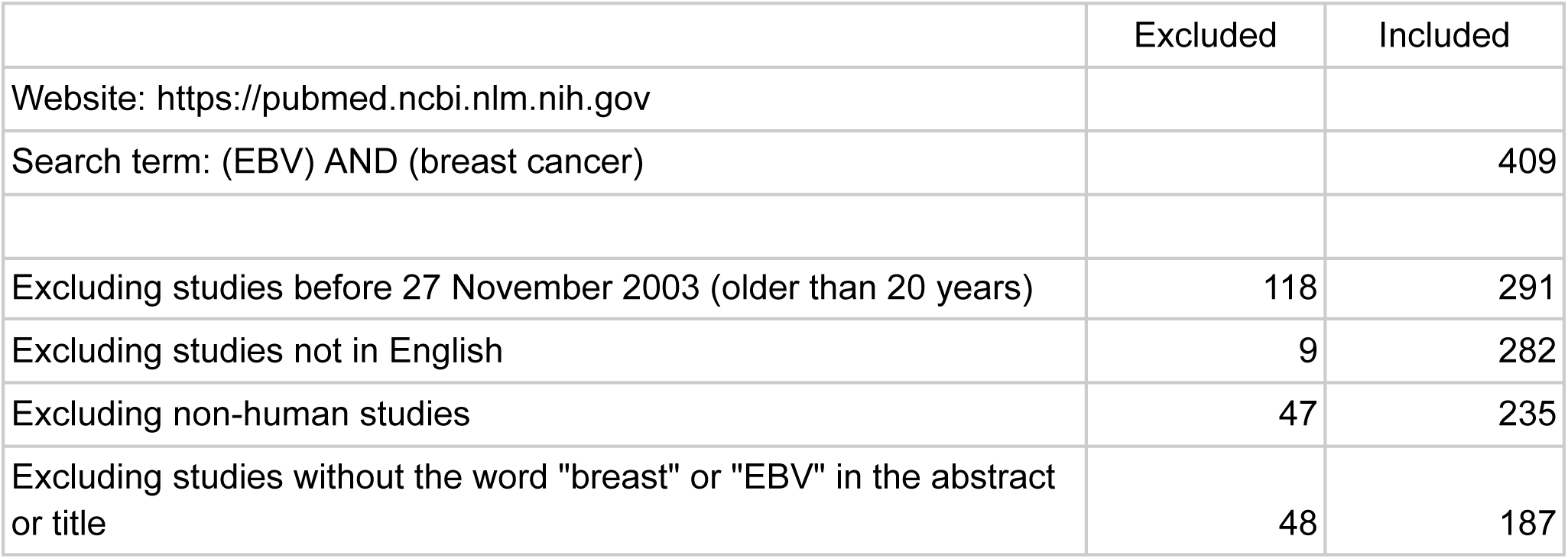
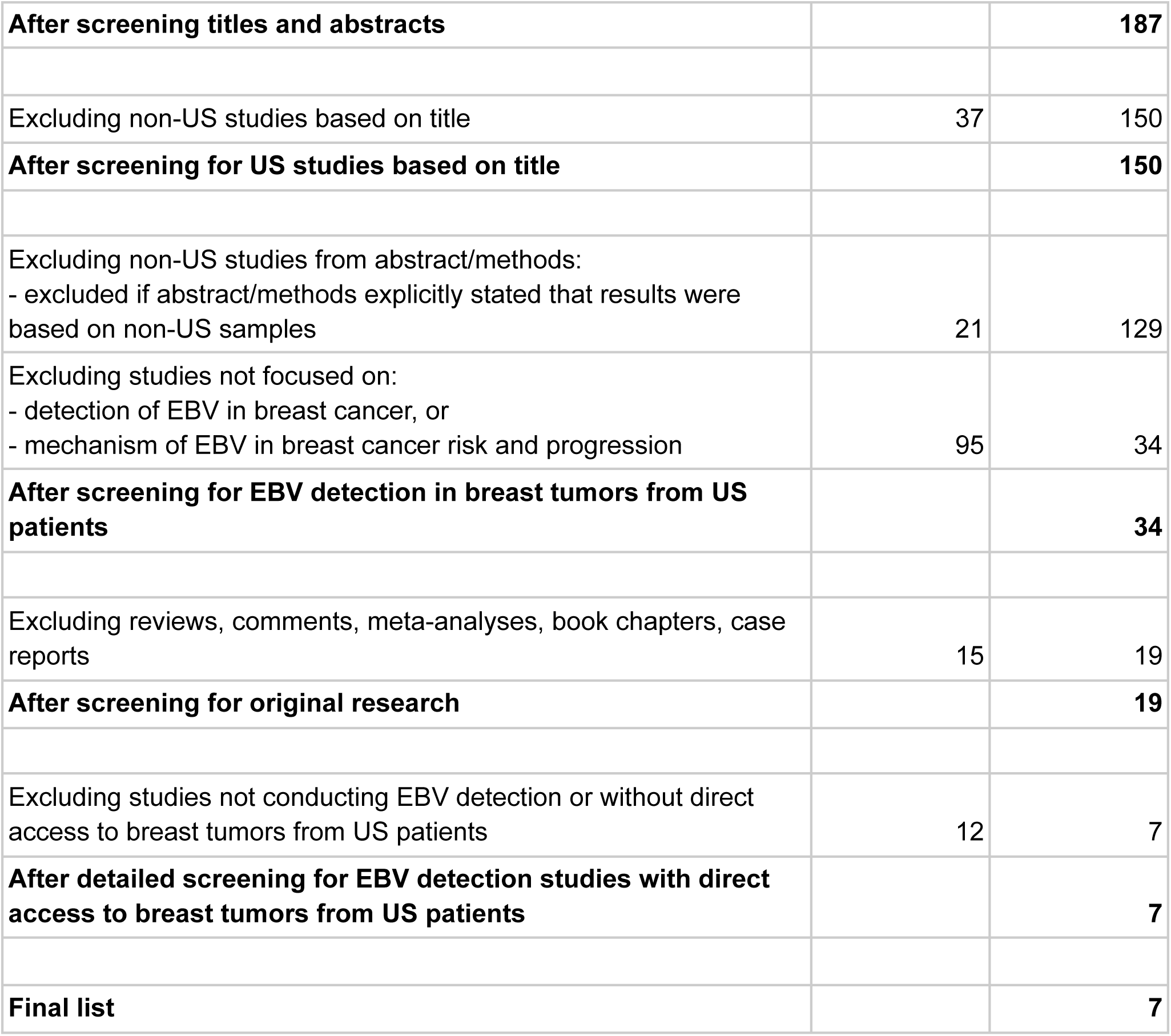

|  | Excluded | Included |
| --- | --- | --- |
| Website: https://pubmed.ncbi.nlm.nih.gov |  |  |
| Search term: (EBV) AND (breast cancer) |  | 409 |
| Excluding studies before 27 November 2003 (older than 20 years) | 118 | 291 |
| Excluding studies not in English | 9 | 282 |
| Excluding non-human studies | 47 | 235 |
| Excluding studies without the word "breast" or "EBV" in the abstract or title | 48 | 187 |
| **After screening titles and abstracts** |  | **187** |
| Excluding non-US studies based on title | 37 | 150 |
| **After screening for US studies based on title** |  | **150** |
| Excluding non-US studies from abstract/methods:<br>- excluded if abstract/methods explicitly stated that results were based on non-US samples | 21 | 129 |
| Excluding studies not focused on:<br>- detection of EBV in breast cancer, or<br>- mechanism of EBV in breast cancer risk and progression | 95 | 34 |
| **After screening for EBV detection in breast tumors from US patients** |  | **34** |
| Excluding reviews, comments, meta-analyses, book chapters, case reports | 15 | 19 |
| **After screening for original research** |  | **19** |
| Excluding studies not conducting EBV detection or without direct access to breast tumors from US patients | 12 | 7 |
| **After detailed screening for EBV detection studies with direct access to breast tumors from US patients** |  | **7** |
| **Final list** |  | **7** |

### Reference Genome Analysis

Given that not all studies published their reference genomes, we analyzed the complete EBV genomes available prior to certain designated study publication years, 2006 and 2012, in an effort to determine whether EBV strains derived from adenocarcinoma cells were used.

### Pre-2006 EBV Genomes

Because 5/7 (71%) studies were published before 2006 or used EBV sequences from a study published before 2006, we first identified and analyzed the complete EBV genomes available before 2006. For this purpose, we queried NCBI GenBank for viral genomic DNA related to EBV (taxonomy ID 10376), published between 1970 and 2006, of reasonable length for complete EBV genomes, and excluded entries labeled with terms indicating non-complete genomes.

To reproduce our results:

1. Visit https://www.ncbi.nlm.nih.gov/nuccore
2. Use this search query:

txid10376[Organism:noexp] AND (viruses[filter] AND biomol_genomic[PROP] AND ddbj_embl_genbank[filter] AND (“150000”[SLEN] : “500000”[SLEN])) AND (“1970/01/01”[PDAT] : “2006/01/01”[PDAT]) NOT (“partial genome”[Title] OR “nearly complete genome”[Title])

### Pre-2012 EBV Genomes

Because one study was published during or after 2006 but before 2012, we also identified and analyzed the complete EBV genomes available before 2012. For this purpose, we used the same query for pre-2006 EBV genomes but updated the date filters.

To reproduce our results:

1. Visit https://www.ncbi.nlm.nih.gov/nuccore
2. Use this search query:

txid10376[Organism:noexp] AND (viruses[filter] AND biomol_genomic[PROP] AND ddbj_embl_genbank[filter] AND (“150000”[SLEN] : “500000”[SLEN])) AND (“1970/01/01”[PDAT] : “2012/01/01”[PDAT]) NOT (“partial genome”[Title] OR “nearly complete genome”[Title])

### Sequence Analysis

The analysis focused on identifying and reporting the number of complete EBV genomes that showed exact matches to the sequences utilized in each study. This was achieved by comparing the complete genome sequences in the dataset against the reference sequences from the studies.

To accomplish this, we searched for EBV Genomes available in 2024 as outlined below and downloaded all the complete genomes in FASTA format.

Next, we compiled the EBV sequences used in each study’s detection protocol.

One issue was that some studies did not label sequence orientation. To avoid overstating mismatches and to adopt the same algorithm for all studies, we analyzed both the original sequence from the study and its reverse complement.

We executed the algorithm below:

1. For a given study sequence, remove white space and label the sequence, “Original.”
2. Generate the reverse complement of “Original” and label this sequence, “RC.”
3. For “Original,” identify the number of EBV genomes with exact matches.
4. Repeat step 3 for “RC.”
5. For each “Original” and “RC” pair, report the higher number of exact matches.
6. Repeat for all sequences.

Python version 3.12.5 and Biopython version 1.81 were used. Code is available at https://github.com/HotpotBio.

### 2024 EBV Genomes

To identify complete EBV genomes available in 2024, we used the same query for pre-2012 EBV genomes but removed the date filters.

To reproduce our results:

1. Visit https://www.ncbi.nlm.nih.gov/nuccore.
2. Use this search query:

txid10376[Organism:noexp] AND (viruses[filter] AND biomol_genomic[PROP] AND ddbj_embl_genbank[filter] AND (“150000”[SLEN] : “500000”[SLEN])) NOT (“partial genome”[Title] OR “nearly complete genome”[Title])

## Results

### Detection Protocols

0/7 (0%) studies utilized sequences of EBV strains extracted from adenocarcinoma cells.

7/7 (100%) studies employed either the Raji or Namalwa cell line as positive controls. Both are derived from EBV-associated Burkitt lymphoma in African patients.

6/7 (86%) of studies either used the B95-8 strain as the reference genome, were published before 2006, or reused sequences from a pre-2006 study. Because the first EBV genome derived from a patient with carcinoma, the GD1 strain, was released in 2006, prior studies necessarily featured lymphoma-derived strains. The one study published after GD1 did not report its reference genome but selected Raji for its positive control. Even if this study used GD1, the reference genome would represent an isolate extracted from saliva and not cancer cells, much less adenocarcinoma cells.

Collectively, this data demonstrates that none of the studies included carcinoma-derived strains, much less adenocarcinoma-derived strains.

**Table 1:** Summary of reference genomes and positive controls. False negative results are more likely without suitable reference genomes and positive controls. EBV and HPV data suggests that utilizing adenocarcinoma-derived strains would mitigate the risk of false negatives when detecting EBV in breast cancer cells. However, all studies either used the B95-8 or Raji strain as reference genomes; the Namalwa, Raji, or Daudi cell lines as positive controls; or sequences available before the first carcinoma genome was published in 2006. This necessarily means no studies used adenocarcinoma-derived strains. - This indicates that the data was not available or not possible to deduce. * This study reused sequences from a 2005 study, so we considered the publication date as equivalent to pre-2006 within the context of reference genomes.(100)

| Study | Authors, Year Published | Reference Genome | Positive Control | Published Before 1st Carcinoma-Derived Genome? |
| --- | --- | --- | --- | --- |
| Absence of the Epstein-Barr virus genome in breast cancer-derived cell line(98) | Speck P. et al, 2003 | - | Namalwa | Yes |
| Lytic viral replication as a contributor to the detection of Epstein-Barr virus in breast cancer(99) | Huang J., 2003 | B95-8, Raji | Namalwa | Yes |
| Real-time PCR measures Epstein-Barr Virus DNA in archival breast adenocarcinomas(100) | Thorne L., 2005 | - | Namalwa, Raji | Yes |
| Lack of association between EBV and breast carcinoma(101) | Perrijoue J., 2005 | - | Raji | Yes |
| Analysis of Epstein-Barr virus reservoirs in paired blood and breast cancer primary biopsy specimens by real time PCR(102) | Perkins S., 2006 | B95-8 | Namalwa, Daudi | - |
| Epstein-Barr virus is seldom found in mammary epithelium of breast cancer tissue using in situ molecular methods(103) | Baltzell K., 2012 | - | Raji | No |
| Variation in risk and outcomes of Epstein-Barr virus-associated breast cancer by epidemiologic characteristics and virus | Glaser S., 2017 | - | Namalwa | Yes* |
| detection strategies: an exploratory study(104) |  |  |  |  |

### Sequence Analysis

7/7 (100%) studies were published before 2012 or used sequences from a study published before 2012. Only ten complete EBV genomes were available before 2012, representing 2% of the 512 complete EBV genomes available in 2024. This means the detection assay sequences in these studies were selected before 98% of the NCBI GenBank entries became available.

5/7 (71%) studies were either published before 2006 or used sequences from a study published before 2006. Only four complete EBV genomes were available before 2006, representing less than 1% of the genomes available in 2024.

5/7 (71%) studies used at least one sequence that did not exactly match more than 50% of EBV genomes. Notably, every study contained at least one sequence that did not exactly match more than 25% of EBV genomes.

**Table 2** lists the assay sequence for each study with the most number of EBV genomes that did not exactly match. See **Supplementary Table 1** for full results of this analysis.

**Table 2:**
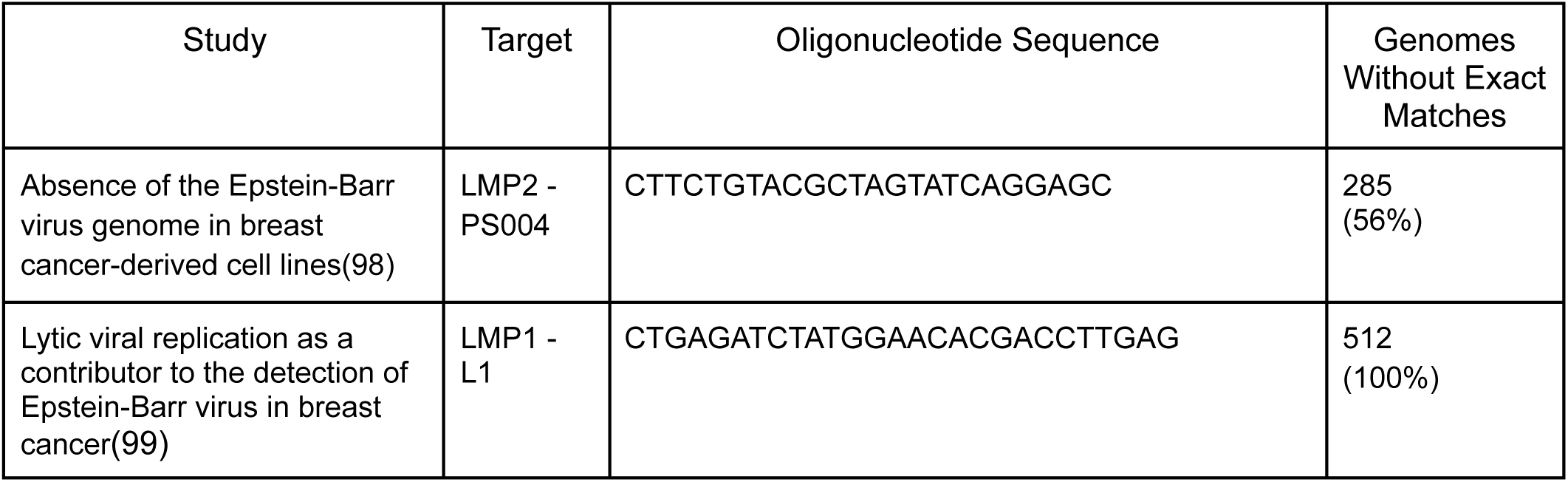
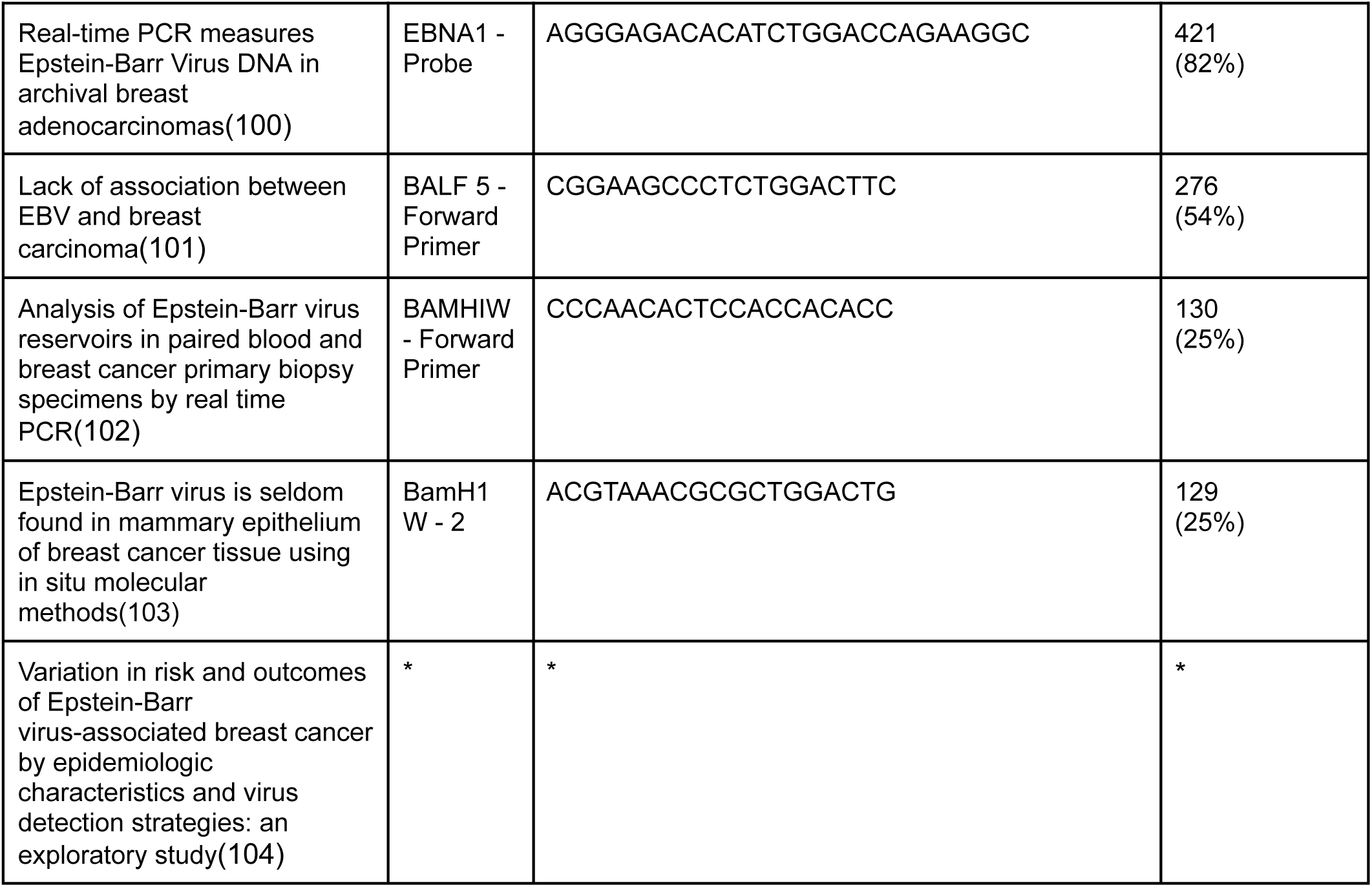
Summary of Genome Exact Match Analysis. 5/7 (71%) studies used at least one sequence that did not exactly match more than 50% of EBV genomes. Notably, every study contained at least one sequence that did not exactly match more than 25% of EBV genomes. These sequences may be less widely conserved than believed and potentially suboptimal for detecting novel variants. Percentages are based on the 512 complete genomes available in May 2024 in NCBI GenBank. This table highlights the sequence for each study with the most number of EBV genomes that did not exactly match. See **Supplementary** Table 1 for full results of the genome-sequence analysis. * This study reused sequences from a 2005 study.(100)

### Biomolecule Integrity

0/7 (0%) of studies reported RIN values despite 5/7 (71%) of studies targeting non-coding RNA in their detection protocols.

## Discussion

This study aimed to reevaluate the association between EBV and breast cancer by analyzing the methodologies and findings of previous research in the context of current genomic data and advanced detection techniques. The results suggest limitations in prior detection protocols. Our analysis highlights several issues, including the potential for false negatives due to reliance on lymphoma-derived strains, overestimating sequence conservation of detection assay sequences, and inadequate addressing of biomolecule integrity, all of which raise concerns about the current understanding of EBV’s role in breast cancer.

### Lymphoma-biased Detection Protocols

False negative results in detection assays are more likely without appropriate reference genomes and positive controls.

All the selected studies relied on the B95-8 and Raji strains, Namalwa, Raji, and Daudi cell lines, or non-carcinoma-derived sequences. This indicates that every one of their detection protocols exclusively relied on lymphoma-derived strains that may not be representative of the strains infective toward breast cancer cells. Given that breast cancers are predominantly adenocarcinomas, utilizing strains extracted from adenocarcinoma cells as reference genomes and positive controls would potentially mitigate the risk of false negatives.

There is insufficient evidence to justify the use of lymphoma-derived strains in these detection protocols. In fact, the current knowledge on EBV and HPV challenges this assumption since different genotypes vary in their pathogenic potential and cell tropism. Notably, only about 5% of HPV genotypes are carcinogenic and a certain EBV strain, namely BALF2_CCT, has demonstrated a 11-times greater risk of NPC due partially to preferential infection of epithelial cells. In fact, even within adenocarcinoma cells it has been demonstrated that the presence of EBV depends on the grade of differentiation, as EBV-detection rates were 8.3% for well-differentiated, 47.2% for moderately differentiated, and 27.6% for poorly differentiated lung adenocarcinomas.(105)

Moreover, the cellular origin of these strains were not only non-glandular and non-epithelial, but in fact non-oncogenic in some cases. The concurrent presence of multiple EBV strains within the same individual suggests that the EBV isolates in cancer cells may differ from those in the non-cancer cells. This underscores the need to restrict reference strains of detection assays to those derived from tumor cells, avoiding ones obtained from saliva and non-tumor cells such as GD1.

Even when utilizing reference genomes and positive controls derived from adenocarcinoma cells, appropriate measures to consider divergent mutations may be necessary to detect novel variants specific to breast glandular cells.

### Unvalidated Sequence Conservation

Kim et al. (2017) observed that “strain variation exists in the EBV genome … such that primers for PCR amplification should target highly conserved sequences in the genome to enable reliable quantification of EBV DNA across different EBV strains/isolates/regional variants.”(106)

However, the selected studies may have potentially overestimated sequence conservation.

Assumptions about EBV sequence conservation merit revalidation given the mounting literature demonstrating greater genomic variation than previously believed, particularly in key coding and non-coding regions. Our analysis reinforces this trend, reporting that 5/7 (71%) studies used at least one assay sequence that did not exactly match more than 50% of EBV genomes. Moreover, 98% of these GenBank entries became available after assay sequences were selected. Because every study targeted multiple sequences, mismatches in one sequence does not invalidate the detection protocol. However, it highlights the need to verify sequence conservation and ensure that protocols reflect the latest genomic data to minimize the risk of overlooking strains potentially tropic to breast cancer cells.

Moreover, while one mismatch is acceptable under the right conditions, the current knowledge on oncoviruses indicates that detecting viruses in adenocarcinomas may require greater sensitivity and specificity. Giannella et al. report that viral DNA may present in lower copy numbers in glandular cells compared to squamous cells and that even single mismatches may trigger false negatives. While this has been demonstrated only in HPV, it may also apply to EBV associated-carcinomas and underscores the need to use the most conserved sequences.(107) Furthermore, one mismatch may lead to lower-reported copies of EBV, which could underestimate the viral load.

### Unverified Biomolecule Integrity

While the gold standard for EBV detection entails EBER in-situ hybridization (EBER-ISH), low RIN scores may compromise the accuracy of such RNA-based assays. Given the higher susceptibility of RNA for degradation due to its single-stranded nature and ubiquitous presence of RNases, ensuring high-quality RNA is paramount. RNA degradation not only reduces the number of EBER molecules but may also impair probe specificity by fragmenting RNA and altering secondary structures.

5/7 (71%) of studies targeted non-coding RNA, but none of them validated biomolecule quality in tissue samples. While some of these studies used GAPDH detection as a surrogate for RNA integrity, RIN scores are a more accurate method of determining RNA integrity. And without adequate validation of biomolecule integrity, it is difficult to determine if the negative results stemmed from sample degradation or absence of EBV.

## Conclusion

In conclusion, we observed a potential risk of failing to detect novel EBV strains in breast tumors based on multiple considerations: lymphoma-biased detection protocols, unvalidated sequence conservation, and unverified biomarker quality. This implies that the current understanding of association between EBV and breast cancer may remain not only incomplete but possibly incorrect.

Given the complex heterogeneity of breast cancer with diverse molecular and histological subtypes such as BRCA1, BRCA2, and TNBC, EBV association with all subtypes is improbable. Nonetheless, should breast cancer mirror gastric carcinoma and reveal viral influence in certain subtypes at certain stages -- initiation, progression, or therapeutic-resistance -- these insights could enable targeted therapies and screening programs. Therefore, we urge renewed investigations into the association between EBV and breast cancer based on gold standard detection protocols that account for novel strains and breast glandular tropism.

## Limitations

Firstly, it is likely that EBV does not influence breast cancer pathogenesis or therapeutic response in American patients. Even if an association is found, the complex heterogeneity of breast cancer with diverse molecular and histological subtypes such as BRCA1, BRCA2, and TNBC suggests that any association may be limited in scope. Moreover, the relationship may be non-causative or immaterial to pathogenesis.

Secondly, reports of EBV association with breast cancer may stem from false positive results due to materials contaminated with EBV, cross-reaction with other markers, or inappropriate detection methods.

Thirdly, it is possible that pre-2006 or pre-2012 genome sequences from lymphoma-derived strains are suitable for detecting oncogenic variants in breast adenocarcinomas. Moreover, HPV and EBV are distinct viruses, and parallels between them may not apply.

## Data Availability

All data underlying our research can be found or reproduced in these sections: **Materials and Methods** and **Supplementary Materials**.

Instructions for accessing the code supporting the genome-sequence analysis can be found under **Materials and Methods**.

## Conflicts of Interest Statement

The authors have no conflicts of interest to declare.

## Funding

This work was supported by Hotpot.ai.

## Acknowledgements

The funder, Hotpot.ai, through its founder and senior author C.H., played a role in the design of the study; the collection, analysis, and interpretation of the data; the writing of the manuscript; and the decision to submit the manuscript for publication.

## Supplementary Material

**Supplementary Table 1:**
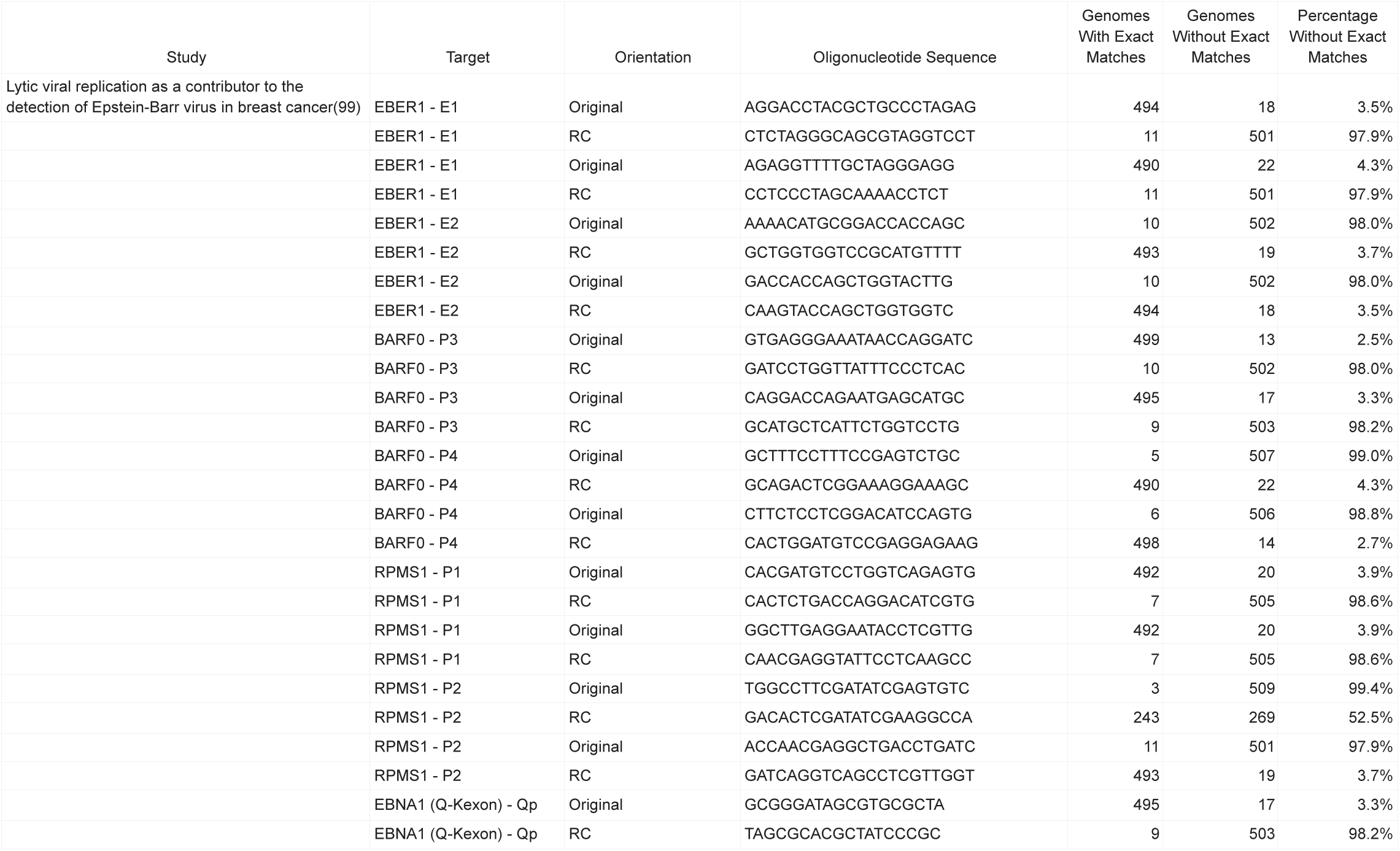
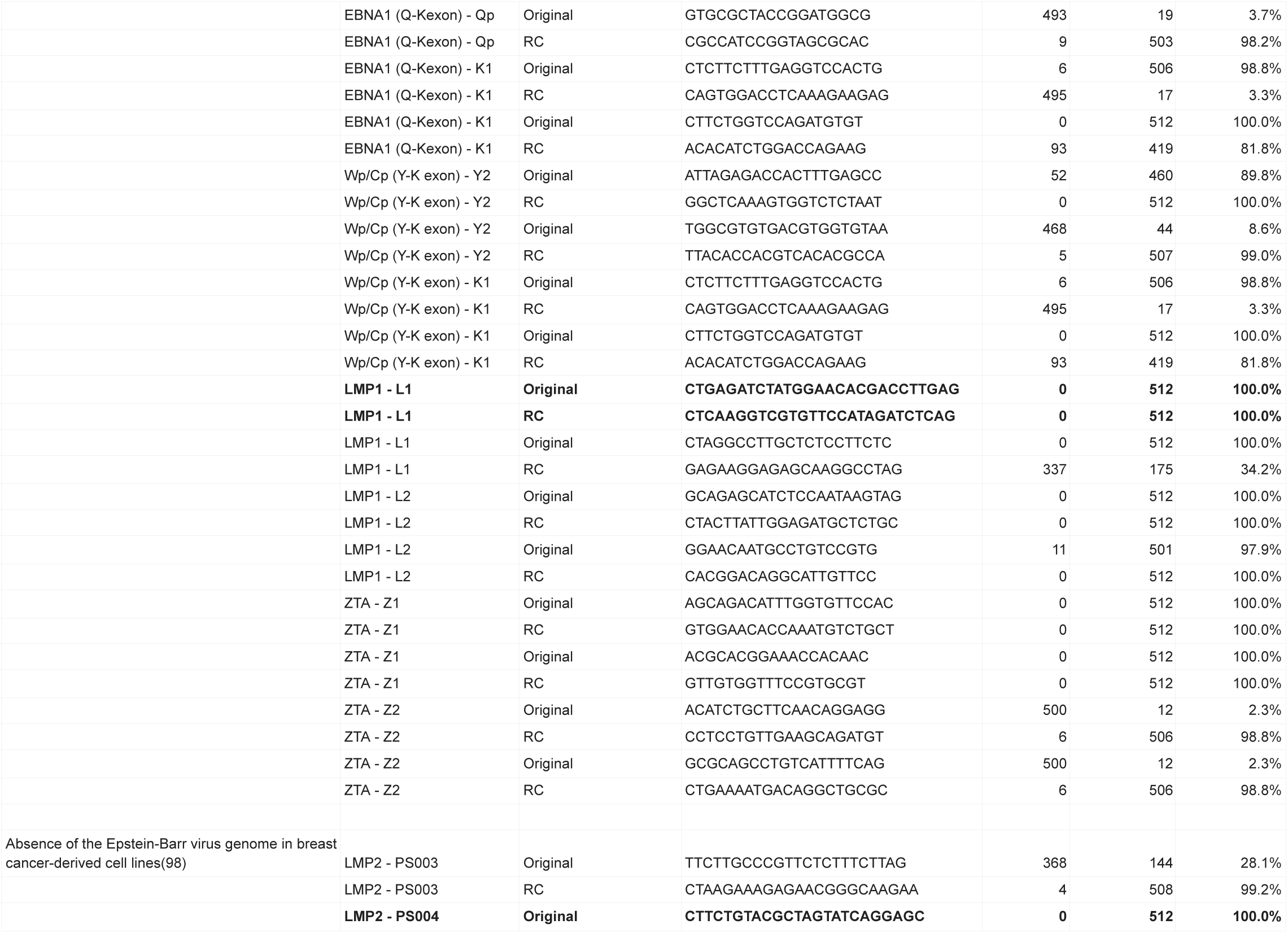
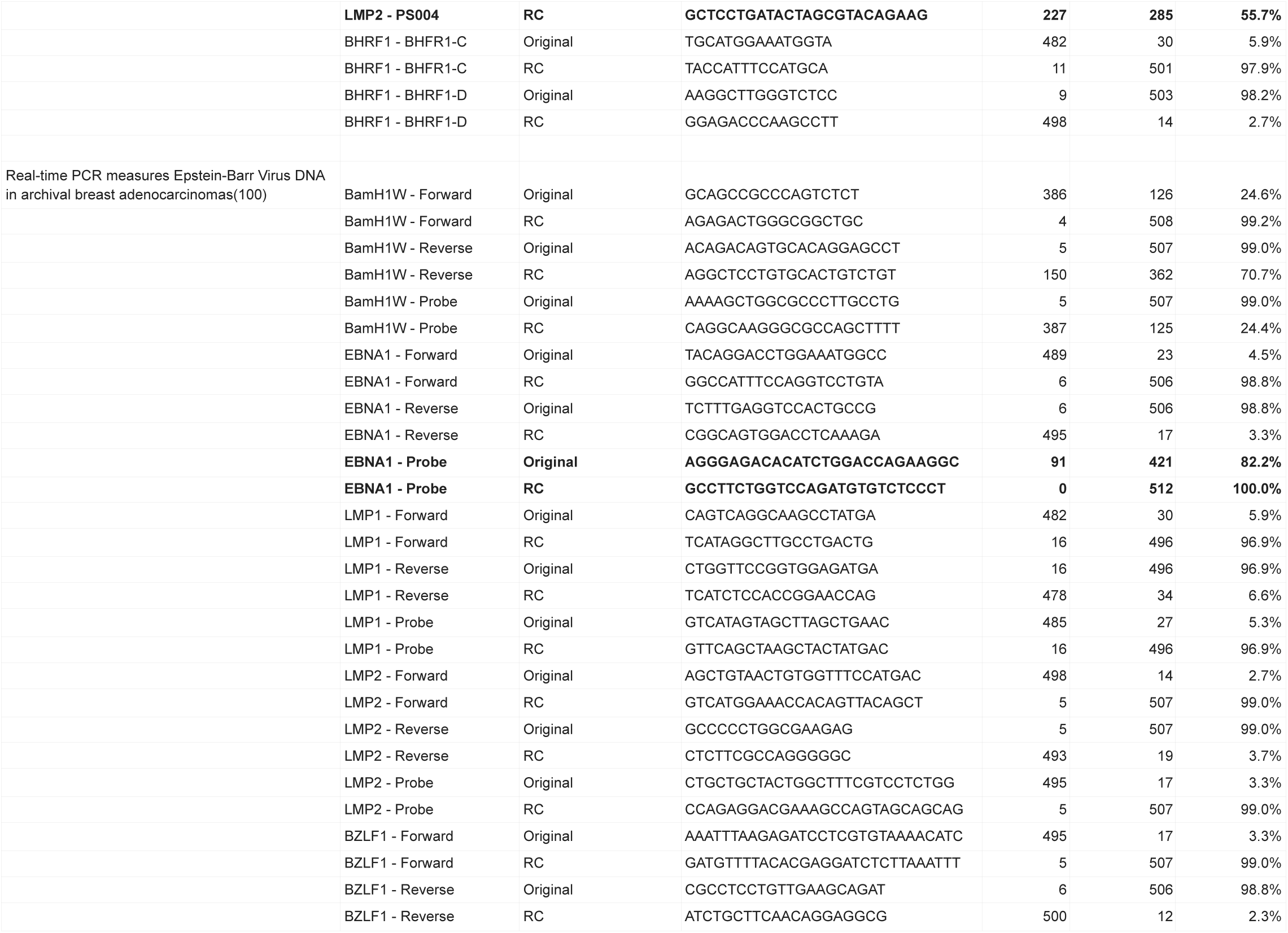
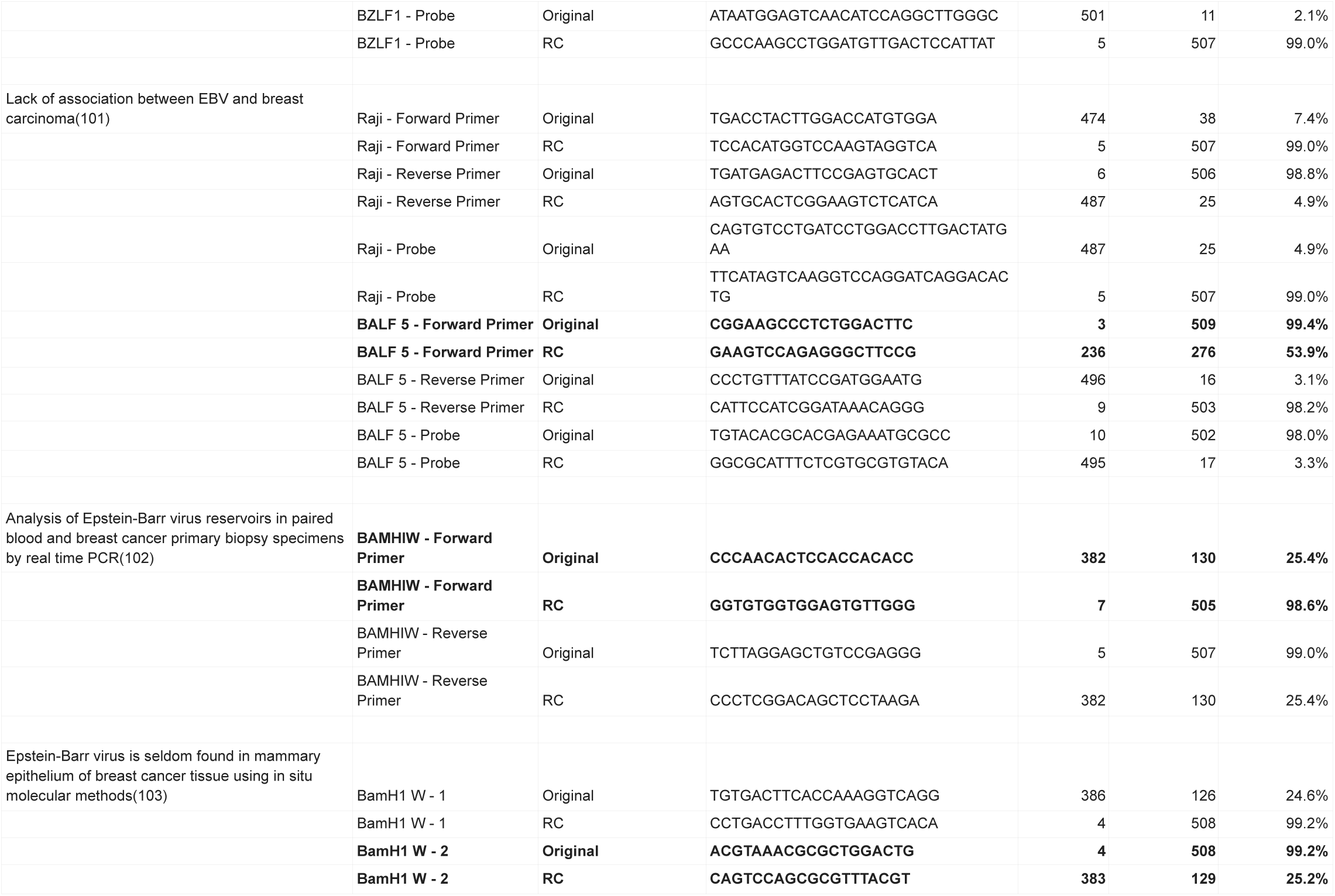
Results of Genome Exact Match Analysis. This table compiles the assay sequences for each study and analyzes the number of EBV genomes with exact matches. An orientation of “Original” reflects the original sequence listed in the study. “RC” represents the reverse complement of “Original.” We test both orientations because some studies did not report orientation, and this approach ensures a consistent method across studies. For each “Original/RC” pair, we report the higher number of genomes with exact matches. Within each study, the bolded pair designates the sequence with the fewest number of exact matches. Percentages are based on the 512 complete genomes available in May 2024 in NCBI GenBank.

## Notes

### Competing Interest Statement

The authors have declared no competing interest.

### Summary of Updates

Added "Objectives" section to better guide readers through the manuscript

## References

1. Plummer M, De Martel C, Vignat J, Ferlay J, Bray F, Franceschi S. Global burden of cancers attributable to infections in 2012: a synthetic analysis. Lancet Glob Health. 2016 Sep;4(9):e609–16.

2. Zapatka M, Borozan I, Brewer DS, Iskar M, Grundhoff A, Alawi M, et al. The landscape of viral associations in human cancers. Nat Genet. 2020 Mar 2;52(3):320–30.

3. Rossi NM, Dai J, Xie Y, Lou H, Boland JF, Yeager M, et al. Extrachromosomal Amplification of Human Papillomavirus Episomes as a Mechanism of Cervical Carcinogenesis [Internet]. 2021 [cited 2024 Jun 26]. Available from: http://biorxiv.org/lookup/doi/10.1101/2021.10.22.465367

4. Aninye IO, Berry-Lawhorn JM, Blumenthal P, Felder T, Jay N, Merrill J, et al. Gaps and Opportunities to Improve Prevention of Human Papillomavirus-Related Cancers. J Womens Health. 2021 Dec 1;30(12):1667–72.

5. Kennedy G, Komano J, Sugden B. Epstein-Barr virus provides a survival factor to Burkitt’s lymphomas. Proc Natl Acad Sci. 2003 Nov 25;100(24):14269–74.

6. Sausen DG, Basith A, Muqeemuddin S. EBV and Lymphomagenesis. Cancers. 2023 Apr 4;15(7):2133.

7. Sun K, Jia K, Lv H, Wang SQ, Wu Y, Lei H, et al. EBV-Positive Gastric Cancer: Current Knowledge and Future Perspectives. Front Oncol. 2020 Dec 14;10:583463.

8. Jary A, Veyri M, Gothland A, Leducq V, Calvez V, Marcelin AG. Kaposi’s Sarcoma-Associated Herpesvirus, the Etiological Agent of All Epidemiological Forms of Kaposi’s Sarcoma. Cancers. 2021 Dec 9;13(24):6208.

9. Yang T, You C, Meng S, Lai Z, Ai W, Zhang J. EBV Infection and Its Regulated Metabolic Reprogramming in Nasopharyngeal Tumorigenesis. Front Cell Infect Microbiol. 2022 Jul 1;12:935205.

10. Yuan L, Li S, Chen Q, Xia T, Luo D, Li L, et al. EBV infection-induced GPX4 promotes chemoresistance and tumor progression in nasopharyngeal carcinoma. Cell Death Differ. 2022 Aug;29(8):1513–27.

11. Hau PM, Lung HL, Wu M, Tsang CM, Wong KL, Mak NK, et al. Targeting Epstein-Barr Virus in Nasopharyngeal Carcinoma. Front Oncol. 2020 May 14;10:600.

12. Kirtane K, St John M, Fuentes-Bayne H, Patel SP, Mardiros A, Xu H, et al. Genomic Immune Evasion: Diagnostic and Therapeutic Opportunities in Head and Neck Squamous Cell Carcinomas. J Clin Med. 2022 Dec 7;11(24):7259.

13. Suresh A, Dhanasekaran R. Implications of genetic heterogeneity in hepatocellular cancer. Adv Cancer Res. 2022;156:103–35.

14. Ruf IK, Rhyne PW, Yang H, Borza CM, Hutt-Fletcher LM, Cleveland JL, et al. Epstein-Barr Virus Regulates c-MYC, Apoptosis, and Tumorigenicity in Burkitt Lymphoma. Mol Cell Biol. 1999 Mar 1;19(3):1651–60.

15. Prusinkiewicz MA, Mymryk JS. Metabolic Control by DNA Tumor Virus-Encoded Proteins. Pathogens. 2021 May 6;10(5):560.

16. Beer S, Wange LE, Zhang X, Kuklik-Roos C, Enard W, Hammerschmidt W, et al. EBNA2-EBF1 complexes promote MYC expression and metabolic processes driving S-phase progression of Epstein-Barr virus–infected B cells. Proc Natl Acad Sci. 2022 Jul 26;119(30):e2200512119.

17. Wang LW, Shen H, Nobre L, Ersing I, Paulo JA, Trudeau S, et al. Epstein-Barr-Virus-Induced One-Carbon Metabolism Drives B Cell Transformation. Cell Metab. 2019 Sep;30(3):539–555.e11.

18. Aloni-Grinstein R, Charni-Natan M, Solomon H, Rotter V. p53 and the Viral Connection: Back into the Future ‡. Cancers. 2018 Jun 4;10(6):178.

19. Saridakis V, Sheng Y, Sarkari F, Holowaty MN, Shire K, Nguyen T, et al. Structure of the p53 Binding Domain of HAUSP/USP7 Bound to Epstein-Barr Nuclear Antigen 1. Mol Cell. 2005 Apr;18(1):25–36.

20. Schönrich G, Raftery MJ. The PD-1/PD-L1 Axis and Virus Infections: A Delicate Balance. Front Cell Infect Microbiol. 2019 Jun 13;9:207.

21. Sasaki S, Nishikawa J, Sakai K, Iizasa H, Yoshiyama H, Yanagihara M, et al. EBV-associated gastric cancer evades T-cell immunity by PD-1/PD-L1 interactions. Gastric Cancer. 2019 May;22(3):486–96.

22. Wang J, Ge J, Wang Y, Xiong F, Guo J, Jiang X, et al. EBV miRNAs BART11 and BART17-3p promote immune escape through the enhancer-mediated transcription of PD-L1. Nat Commun. 2022 Feb 14;13(1):866.

23. Bi XW, Wang H, Zhang WW, Xia Z, Zhang Y jing, Wang L. PD-L1 Is up-Regulated By EBV-Driven LMP1 through NF-κb Pathway and Correlates with Poor Prognosis in Natural Killer/T-Cell Lymphoma. Blood. 2016 Dec 2;128(22):4134–4134.

24. Nakano H, Saito M, Nakajima S, Saito K, Nakayama Y, Kase K, et al. PD-L1 overexpression in EBV-positive gastric cancer is caused by unique genomic or epigenomic mechanisms. Sci Rep. 2021 Jan 21;11(1):1982.

25. Yanagi Y, Hara Y, Mabuchi S, Watanabe T, Sato Y, Kimura H, et al. PD-L1 upregulation by lytic induction of Epstein-Barr Virus. Virology. 2022 Mar;568:31–40.

26. Polansky H, Schwab H. How latent viruses cause breast cancer: An explanation based on the microcompetition model. Bosn J Basic Med Sci [Internet]. 2018 Dec 23 [cited 2024 Jun 26]; Available from: http://www.bjbms.org/ojs/index.php/bjbms/article/view/3950

27. Friedenson B. In BRCA1 and BRCA2 breast cancers, chromosome breaks occur near herpes tumor virus sequences [Internet]. 2021 [cited 2024 Jun 26]. Available from: http://biorxiv.org/lookup/doi/10.1101/2021.04.19.440499

28. Lai KM, Lee WL. The roles of epidermal growth factor receptor in viral infections. Growth Factors. 2022 May 4;40(1–2):46–72.

29. Ho J, Moyes DL, Tavassoli M, Naglik JR. The Role of ErbB Receptors in Infection. Trends Microbiol. 2017 Nov;25(11):942–52.

30. Pekkonen P, Järviluoma A, Zinovkina N, Cvrljevic A, Prakash S, Westermarck J, et al. KSHV viral cyclin interferes with T-cell development and induces lymphoma through Cdk6 and Notch activation in vivo. Cell Cycle. 2014 Dec;13(23):3670–84.

31. Komatsu H, Imadome KI, Shibayama H, Yada T, Yamada M, Yamamoto K, et al. STAT3 Is Activated By EBV in T or NK Cells Leading to Development of EBV-T/NK-Lymphoproliferative Disorders. Blood. 2014 Dec 6;124(21):3026–3026.

32. Li JSZ, Abbasi A, Kim DH, Lippman SM, Alexandrov LB, Cleveland DW. Chromosomal fragile site breakage by EBV-encoded EBNA1 at clustered repeats. Nature. 2023 Apr 20;616(7957):504–9.

33. Portis T, Longnecker R. Epstein–Barr virus (EBV) LMP2A mediates B-lymphocyte survival through constitutive activation of the Ras/PI3K/Akt pathway. Oncogene. 2004 Nov 11;23(53):8619–28.

34. Vaysberg M, Lambert SL, Krams SM, Martinez OM. Activation of the JAK/STAT Pathway in Epstein Barr Virus+-Associated Posttransplant Lymphoproliferative Disease: Role of Interferon-γ. Am J Transplant. 2009 Oct;9(10):2292–302.

35. Arbach H, Viglasky V, Lefeu F, Guinebretière JM, Ramirez V, Bride N, et al. Epstein-Barr Virus (EBV) Genome and Expression in Breast Cancer Tissue: Effect of EBV Infection of Breast Cancer Cells on Resistance to Paclitaxel (Taxol). J Virol. 2006 Jan 15;80(2):845–53.

36. Rüeger S, Hammer C, Loetscher A, McLaren PJ, Lawless D, Naret O, et al. The influence of human genetic variation on Epstein–Barr virus sequence diversity. Sci Rep. 2021 Feb 25;11(1):4586.

37. Balfour HH, Sifakis F, Sliman JA, Knight JA, Schmeling DO, Thomas W. Age-specific prevalence of Epstein-Barr virus infection among individuals aged 6-19 years in the United States and factors affecting its acquisition. J Infect Dis. 2013 Oct 15;208(8):1286–93.

38. Robinson WH, Steinman L. Epstein-Barr virus and multiple sclerosis. Science. 2022 Jan 21;375(6578):264–5.

39. Soldan SS, Lieberman PM. Epstein–Barr virus and multiple sclerosis. Nat Rev Microbiol. 2023 Jan;21(1):51–64.

40. Proceedings of the IARC Working Group on the Evaluation of Carcinogenic Risks to Humans. Epstein-Barr Virus and Kaposi’s Sarcoma Herpesvirus/Human Herpesvirus 8. Lyon, France, 17-24 June 1997. Iarc Monogr Eval Carcinog Risks Hum. 1997;70:1–492.

41. Farahmand M, Monavari SH, Shoja Z, Ghaffari H, Tavakoli M, Tavakoli A. Epstein–Barr Virus and Risk of Breast Cancer: A Systematic Review and Meta-Analysis. Future Oncol. 2019 Aug 31;15(24):2873–85.

42. Arias-Calvachi C, Blanco R, Calaf GM, Aguayo F. Epstein–Barr Virus Association with Breast Cancer: Evidence and Perspectives. Biology. 2022 May 24;11(6):799.

43. Sinclair AJ, Moalwi MH, Amoaten T. Is EBV Associated with Breast Cancer in Specific Geographic Locations? Cancers. 2021 Feb 16;13(4):819.

44. Gupta I, Ulamec M, Peric-Balja M, Ramic S, Al Moustafa AE, Vranic S, et al. Presence of high-risk HPVs, EBV, and MMTV in human triple-negative breast cancer. Hum Vaccines Immunother. 2021 Nov 2;17(11):4457–66.

45. Mekrazi S, Kallel I, Jamai D, Yengui M, Khabir A, Gdoura R. Epstein-Barr virus in breast carcinoma and in triple negative cases impact on clinical outcomes. Pathol Res Pract. 2023 May;245:154484.

46. Hu H, Luo ML, Desmedt C, Nabavi S, Yadegarynia S, Hong A, et al. Epstein-Barr Virus Infection of Mammary Epithelial Cells Promotes Malignant Transformation. EBioMedicine. 2016 Jul;9:148–60.

47. Farrell PJ. Epstein–Barr Virus and Cancer. Annu Rev Pathol Mech Dis. 2019 Jan 24;14(1):29–53.

48. Nanbo A, Inoue K, Adachi-Takasawa K, Takada K. Epstein–Barr virus RNA confers resistance to interferon-α-induced apoptosis in Burkitt’s lymphoma. EMBO J. 2002 Mar 1;21(5):954.

49. Kitagawa N, Goto M, Kurozumi K, Maruo S, Fukayama M, Naoe T, et al. Epstein–Barr virus-encoded poly(A)– RNA supports Burkitt’s lymphoma growth through interleukin-10 induction. EMBO J. 2000 Dec 15;19(24):6742.

50. Farrell PJ, White RE. Do Epstein–Barr Virus Mutations and Natural Genome Sequence Variations Contribute to Disease? Biomolecules. 2021 Dec 23;12(1):17.

51. Borozan I, Zapatka M, Frappier L, Ferretti V. Analysis of Epstein-Barr Virus Genomes and Expression Profiles in Gastric Adenocarcinoma. Longnecker RM, editor. J Virol. 2018 Jan 15;92(2):e01239–17.

52. Wang HY, Sun L, Li P, Liu W, Zhang ZG, Luo B. Sequence Variations of Epstein-Barr Virus-Encoded Small Noncoding RNA and Latent Membrane Protein 1 in Hematologic Tumors in Northern China. Intervirology. 2021;64(2):69–80.

53. Young LS. A novel Epstein-Barr virus subtype associated with nasopharyngeal carcinoma found in South China. Cancer Commun. 2020 Jan;40(1):60–2.

54. Kwok H, Chiang A. From Conventional to Next Generation Sequencing of Epstein-Barr Virus Genomes. Viruses. 2016 Feb 24;8(3):60.

55. Young LS, Yap LF, Murray PG. Epstein-Barr virus: more than 50 years old and still providing surprises. Nat Rev Cancer. 2016 Dec;16(12):789–802.

56. Tu C, Zeng Z, Qi P, Li X, Guo C, Xiong F, et al. Identification of genomic alterations in nasopharyngeal carcinoma and nasopharyngeal carcinoma-derived Epstein–Barr virus by whole-genome sequencing. Carcinogenesis. 2018 Dec 31;39(12):1517–28.

57. Fitzpatrick MB, Hahn Z, Mandishora RSD, Dao J, Weber J, Huang C, et al. Whole-Genome Analysis of Cervical Human Papillomavirus Type 35 from rural Zimbabwean Women. Sci Rep. 2020 Apr 24;10(1):7001.

58. Kanda T, Yajima M, Ikuta K. Epstein-Barr virus strain variation and cancer. Cancer Sci. 2019 Apr;110(4):1132–9.

59. Palser AL, Grayson NE, White RE, Corton C, Correia S, Ba Abdullah MM, et al. Genome Diversity of Epstein-Barr Virus from Multiple Tumor Types and Normal Infection. Longnecker RM, editor. J Virol. 2015 May 15;89(10):5222–37.

60. Santpere G, Darre F, Blanco S, Alcami A, Villoslada P, Mar Albà M, et al. Genome-Wide Analysis of Wild-Type Epstein–Barr Virus Genomes Derived from Healthy Individuals of the 1000 Genomes Project. Genome Biol Evol. 2014 Apr;6(4):846–60.

61. Tzellos S, Farrell P. Epstein-Barr Virus Sequence Variation—Biology and Disease. Pathogens. 2012 Nov 8;1(2):156–74.

62. Hui KF, Chan TF, Yang W, Shen JJ, Lam KP, Kwok H, et al. High risk Epstein-Barr virus variants characterized by distinct polymorphisms in the EBER locus are strongly associated with nasopharyngeal carcinoma. Int J Cancer. 2019 Jun 15;144(12):3031–42.

63. Wang Y, Zhang X, Chao Y, Jia Y, Xing X, Luo B. New variations of Epstein–Barr virus-encoded small RNA genes in nasopharyngeal carcinomas, gastric carcinomas, and healthy donors in northern China. J Med Virol. 2010 May;82(5):829–36.

64. Wang Y, Ungerleider N, Hoffman BA, Kara M, Farrell PJ, Flemington EK, et al. A Polymorphism in the Epstein-Barr Virus EBER2 Noncoding RNA Drives *In Vivo* Expansion of Latently Infected B Cells. Ackerman ME, editor. mBio. 2022 Jun 28;13(3):e00836–22.

65. Salnikov MY, MacNeil KM, Mymryk JS. The viral etiology of EBV-associated gastric cancers contributes to their unique pathology, clinical outcomes, treatment responses and immune landscape. Front Immunol. 2024 Mar 26;15:1358511.

66. Kondo S, Okuno Y, Murata T, Dochi H, Wakisaka N, Mizokami H, et al. EBV genome variations enhance clinicopathological features of nasopharyngeal carcinoma in a non-endemic region. Cancer Sci. 2022 Jul;113(7):2446–56.

67. Dheekollu J, Malecka K, Wiedmer A, Delecluse HJ, Chiang AKS, Altieri DC, et al. Carcinoma-risk variant of EBNA1 deregulates Epstein-Barr Virus episomal latency. Oncotarget. 2017 Jan 31;8(5):7248–64.

68. Bristol JA, Djavadian R, Albright ER, Coleman CB, Ohashi M, Hayes M, et al. A cancer-associated Epstein-Barr virus BZLF1 promoter variant enhances lytic infection. Flemington EK, editor. PLOS Pathog. 2018 Jul 27;14(7):e1007179.

69. Liu Y, Yang W, Pan Y, Ji J, Lu Z, Ke Y. Genome-wide analysis of Epstein-Barr virus (EBV) isolated from EBV-associated gastric carcinoma (EBVaGC). Oncotarget. 2016 Jan 26;7(4):4903–14.

70. Sousa H, Tavares A, Campos C, Marinho-Dias J, Brito M, Medeiros R, et al. High-Risk human papillomavirus genotype distribution in the Northern region of Portugal: Data from regional cervical cancer screening program. Papillomavirus Res. 2019 Dec;8:100179.

71. Romero-Masters JC, Huebner SM, Ohashi M, Bristol JA, Benner BE, Barlow EA, et al. B cells infected with Type 2 Epstein-Barr virus (EBV) have increased NFATc1/NFATc2 activity and enhanced lytic gene expression in comparison to Type 1 EBV infection. Swaminathan S, editor. PLOS Pathog. 2020 Feb 14;16(2):e1008365.

72. Singh DR, Nelson SE, Pawelski AS, Cantres-Velez JA, Kansra AS, Pauly NP, et al. Type 1 and Type 2 Epstein-Barr viruses induce proliferation, and inhibit differentiation, in infected telomerase-immortalized normal oral keratinocytes. Sample CE, editor. PLOS Pathog. 2022 Oct 3;18(10):e1010868.

73. Rickinson AB, Young LS, Rowe M. Influence of the Epstein-Barr virus nuclear antigen EBNA 2 on the growth phenotype of virus-transformed B cells. J Virol. 1987 May;61(5):1310–7.

74. Xu M, Yao Y, Chen H, Zhang S, Cao SM, Zhang Z, et al. Genome sequencing analysis identifies Epstein-Barr virus subtypes associated with high risk of nasopharyngeal carcinoma. Nat Genet. 2019 Jul;51(7):1131–6.

75. Kwok H, Wu CW, Palser AL, Kellam P, Sham PC, Kwong DLW, et al. Genomic Diversity of Epstein-Barr Virus Genomes Isolated from Primary Nasopharyngeal Carcinoma Biopsy Samples. Hutt-Fletcher LM, editor. J Virol. 2014 Sep 15;88(18):10662–72.

76. Correia S, Bridges R, Wegner F, Venturini C, Palser A, Middeldorp JM, et al. Sequence Variation of Epstein-Barr Virus: Viral Types, Geography, Codon Usage, and Diseases. Longnecker RM, editor. J Virol. 2018 Nov 15;92(22):e01132–18.

77. Xue WQ, Wang TM, Huang JW, Zhang JB, He YQ, Wu ZY, et al. A comprehensive analysis of genetic diversity of EBV reveals potential high-risk subtypes associated with nasopharyngeal carcinoma in China. Virus Evol. 2021 Jan 20;7(1):veab010.

78. Tsai MH, Raykova A, Klinke O, Bernhardt K, Gärtner K, Leung CS, et al. Spontaneous Lytic Replication and Epitheliotropism Define an Epstein-Barr Virus Strain Found in Carcinomas. Cell Rep. 2013 Oct;5(2):458–70.

79. White RE, Rämer PC, Naresh KN, Meixlsperger S, Pinaud L, Rooney C, et al. EBNA3B-deficient EBV promotes B cell lymphomagenesis in humanized mice and is found in human tumors. J Clin Invest. 2012 Apr 2;122(4):1487–502.

80. Li Z, Baccianti F, Delecluse S, Tsai MH, Shumilov A, Cheng X, et al. The Epstein–Barr virus noncoding RNA EBER2 transactivates the UCHL1 deubiquitinase to accelerate cell growth. Proc Natl Acad Sci. 2021 Oct 26;118(43):e2115508118.

81. McCall LI, Siqueira-Neto JL, McKerrow JH. Location, Location, Location: Five Facts about Tissue Tropism and Pathogenesis. Knoll LJ, editor. PLOS Pathog. 2016 May 26;12(5):e1005519.

82. Bu GL, Xie C, Kang YF, Zeng MS, Sun C. How EBV Infects: The Tropism and Underlying Molecular Mechanism for Viral Infection. Viruses. 2022 Oct 27;14(11):2372.

83. NCBI [Internet]. [cited 2024 Jul 25]. Genome. Available from: https://www.ncbi.nlm.nih.gov/datasets/genome/?taxon=10376

84. Zanella L, Riquelme I, Buchegger K, Abanto M, Ili C, Brebi P. A reliable Epstein-Barr Virus classification based on phylogenomic and population analyses. Sci Rep. 2019 Jul 8;9(1):9829.

85. Allen UD, Hu P, Pereira SL, Robinson JL, Paton TA, Beyene J, et al. The genetic diversity of Epstein-Barr virus in the setting of transplantation relative to non-transplant settings: A feasibility study. Pediatr Transplant. 2016 Feb;20(1):124–9.

86. Osunkoya BO. The preservation of Burkitt tumour cells at moderately low temperature. Br J Cancer. 1965 Dec;19(4):749–53.

87. Fryer JF, Heath AB, Wilkinson DE, Minor PD. A collaborative study to establish the 1st WHO International Standard for Epstein–Barr virus for nucleic acid amplification techniques. Biologicals. 2016 Sep;44(5):423–33.

88. Delecluse HJ, Hilsendegen T, Pich D, Zeidler R, Hammerschmidt W. Propagation and recovery of intact, infectious Epstein–Barr virus from prokaryotic to human cells. Proc Natl Acad Sci. 1998 Jul 7;95(14):8245–50.

89. Pulvertaft RJV. CYTOLOGY OF BURKITT’S TUMOUR (AFRICAN LYMPHOMA). The Lancet. 1964 Feb;283(7327):238–40.

90. Young LS, Yao QY, Rooney CM, Sculley TB, Moss DJ, Rupani H, et al. New type B isolates of Epstein-Barr virus from Burkitt’s lymphoma and from normal individuals in endemic areas. J Gen Virol. 1987 Nov;68 (Pt 11):2853–62.

91. Delecluse HJ, Bartnizke S, Hammerschmidt W, Bullerdiek J, Bornkamm GW. Episomal and integrated copies of Epstein-Barr virus coexist in Burkitt lymphoma cell lines. J Virol. 1993 Mar;67(3):1292–9.

92. Henderson A, Ripley S, Heller M, Kieff E. Chromosome site for Epstein-Barr virus DNA in a Burkitt tumor cell line and in lymphocytes growth-transformed in vitro. Proc Natl Acad Sci. 1983 Apr;80(7):1987–91.

93. Pirog EC. Cervical Adenocarcinoma: Diagnosis of Human Papillomavirus–Positive and Human Papillomavirus–Negative Tumors. Arch Pathol Lab Med. 2017 Dec 1;141(12):1653–67.

94. Chow CK, Qin K, Lau LT, Cheung-Hoi Yu A. Significance of a Single-Nucleotide Primer Mismatch in Hepatitis B Virus Real-Time PCR Diagnostic Assays. J Clin Microbiol. 2011 Dec;49(12):4418–9.

95. Walling DM, Brown AL, Etienne W, Keitel WA, Ling PD. Multiple Epstein-Barr Virus Infections in Healthy Individuals. J Virol. 2003 Jun;77(11):6546–50.

96. RNA Integrity Number - an overview | ScienceDirect Topics [Internet]. [cited 2024 Jul 25]. Available from: https://www.sciencedirect.com/topics/biochemistry-genetics-and-molecular-biology/rna-integrity-number

97. Fleige S, Pfaffl MW. RNA integrity and the effect on the real-time qRT-PCR performance. Mol Aspects Med. 2006 Apr;27(2–3):126–39.

98. Speck P, Callen DF, Longnecker R. Absence of the Epstein-Barr virus genome in breast cancer-derived cell lines. J Natl Cancer Inst. 2003 Aug 20;95(16):1253–4; author reply 1254-1255.

99. Huang J, Chen H, Hutt-Fletcher L, Ambinder RF, Hayward SD. Lytic viral replication as a contributor to the detection of Epstein-Barr virus in breast cancer. J Virol. 2003 Dec;77(24):13267–74.

100. Thorne LB, Ryan JL, Elmore SH, Glaser SL, Gulley ML. Real-time PCR measures Epstein-Barr Virus DNA in archival breast adenocarcinomas. Diagn Mol Pathol Am J Surg Pathol Part B. 2005 Mar;14(1):29–33.

101. Perrigoue JG, den Boon JA, Friedl A, Newton MA, Ahlquist P, Sugden B. Lack of association between EBV and breast carcinoma. Cancer Epidemiol Biomark Prev Publ Am Assoc Cancer Res Cosponsored Am Soc Prev Oncol. 2005 Apr;14(4):809–14.

102. Perkins RS, Sahm K, Marando C, Dickson-Witmer D, Pahnke GR, Mitchell M, et al. Analysis of Epstein-Barr virus reservoirs in paired blood and breast cancer primary biopsy specimens by real time PCR. Breast Cancer Res BCR. 2006;8(6):R70.

103. Baltzell K, Buehring GC, Krishnamurthy S, Kuerer H, Shen HM, Sison JD. Epstein-Barr virus is seldom found in mammary epithelium of breast cancer tissue using in situ molecular methods. Breast Cancer Res Treat. 2012 Feb;132(1):267–74.

104. Glaser SL, Canchola AJ, Keegan THM, Clarke CA, Longacre TA, Gulley ML. Variation in risk and outcomes of Epstein-Barr virus-associated breast cancer by epidemiologic characteristics and virus detection strategies: an exploratory study. Cancer Causes Control CCC. 2017 Apr;28(4):273–87.

105. Chen Y, Liu T, Xu Z, Dong M. Association of Epstein-Barr virus (EBV) with lung cancer: meta-analysis. Front Oncol. 2023;13:1177521.

106. Kim KY, Le QT, Yom SS, Pinsky BA, Bratman SV, Ng RHW, et al. Current State of PCR-Based Epstein-Barr Virus DNA Testing for Nasopharyngeal Cancer. JNCI J Natl Cancer Inst. 2017 Mar 14;109(4):djx007.

107. Giannella L, Di Giuseppe J, Delli Carpini G, Grelloni C, Fichera M, Sartini G, et al. HPV-Negative Adenocarcinomas of the Uterine Cervix: From Molecular Characterization to Clinical Implications. Int J Mol Sci. 2022 Nov 30;23(23):15022.

